# Sleep alterations in a mouse model of Spinocerebellar ataxia type 3

**DOI:** 10.1101/2022.07.05.498300

**Authors:** Maria-Efstratia Tsimpanouli, Anjesh Ghimire, Anna J. Barget, Ridge Weston, Henry L. Paulson, Maria do Carmo Costa, Brendon O. Watson

## Abstract

**Background:** Spinocerebellar ataxia type 3 (SCA3) is a neurodegenerative disorder showing progressive neuronal loss in several brain areas and a broad spectrum of motor and non-motor symptoms, including ataxia and altered sleep. While sleep disturbances are known to play pathophysiologic roles in other neurodegenerative disorders, their impact on SCA3 is unknown.

**Objectives:** Using state-of-the art spectrographic measurements, we sought to quantitatively characterize sleep electroencephalography (EEG) in a SCA3 transgenic mouse model with confirmed disease phenotype.

**Methods:** We first measured motor phenotypes in 18–31-week-old homozygous and hemizygous SCA3 YACMJD84.2 mice and non-transgenic wild-type littermate mice during lights-on and lights-off periods. We next implanted electrodes to obtain 12-hour (zeitgeber time 0-12) EEG recordings for three consecutive days when the mice were 26–36 weeks old. We then analyzed EEG-based sleep structure data to quantify differences between homozygous, hemizygous, and wild-type mice.

**Results:** Compared to wild-type littermates, SCA3 homozygous mice display: i) increased duration of rapid-eye movement sleep (REM) and fragmentation in all sleep and wake states; ii) higher beta power oscillations during REM and non-REM (NREM); and iii) additional spectral power band alterations during REM and wake.

**Conclusions:** Our data show that sleep architecture and EEG spectral power are dysregulated in homozygous SCA3 mice, indicating that common sleep-related etiologic factors may underlie mouse and human SCA3 phenotypes.

## INTRODUCTION

Spinocerebellar ataxia type 3 (SCA3), also known as Machado-Joseph disease, is a fatal and incurable dominantly-inherited ataxia with typical adult onset of neurodegenerative symptoms^1^. SCA3 is caused by a CAG repeat expansion in the *ATXN3* gene encoding a polyglutamine stretch in the protein ataxin-3 (ATXN3)^2^, which is ubiquitously expressed throughout the brain and the body^3, 4^. People with SCA3 experience a wide variety of motor impairments and non-motor neurological symptoms, depending on the areas of the central and peripheral nervous system affected by the degenerative process^5–8^. The brain areas that are most frequently affected include the cerebellum, brainstem, cerebral cortex, basal ganglia, thalamus, and midbrain^5, 6^. However, disease onset is determined based on the onset of progressive cerebellar ataxia, which is usually the main SCA3 symptom^9^.

In SCA3, sleep disorders are common and have been identified in up to 60% of patients at early and later stages of disease^9, 10^. In some SCA3 cases, sleep disorders precede ataxia onset by 5-10 years, implying a possible pathophysiologic role for sleep in this disease^11, 12^. The most common sleep disorders in SCA3 are insomnia^13, 14^, excessive daytime sleepiness^15–20^, restless leg syndrome^11, 13, 14, 17, 21–26^, periodic limb movements disorder^15, 17, 23, 27, 28^, REM sleep without atonia^23, 27–29^, REM sleep behavior disorder (RBD)^11–13, 16, 27–31, 32^, obstructive sleep apnea^13, 15, 18^, confusional arousals^15, 28^, and sleep terrors^28, 29^. However, we lack knowledge of sleep structure changes and electrophysiology in SCA3. Such knowledge, combined with details of which neuroanatomical regions are driving those changes, can enable mechanistic studies of the role of sleep in SCA3 progression, as has been implicated in other neurodegenerative diseases such as Huntington (HD), Parkinson (PD), and Alzheimer (AD) diseases. While anatomy-physiology correlates can be studied with spectrographic analysis of electroencephalogram (EEG) data during sleep, only one spectrographic sleep study of SCA1, 2 and 3 patients has been carried out and it only focused on sleep spindles without quantification of other oscillation frequencies^33^.

Neurobiologic mechanistic studies are often most powerfully carried out in mouse models, given the molecular, genetic, and neuroscientific tools available in those models. Mouse models have been instrumental in revealing SCAs’ disease mechanisms. The SCA3 YACMJD84.2 (Q84) transgenic mouse, which expresses the full-length human disease *ATXN3* gene, replicates many aspects of SCA3 including motor impairments and thus is frequently used in preclinical studies of SCA3^34, 35^. Homozygous mice (Q84/Q84) show motor impairments by six weeks of age and hemizygous mice (Q84/WT) show a later-onset mild motor phenotype^35^. Further use of these models in combination with powerful in vivo neurobiological tools can allow us to efficiently understand the impacts of this gene mutation on brain function states in wake and sleep. Despite the overall lack of spectrographic sleep studies in SCA3 mouse models, one previous study analyzing just 30 seconds of EEG identified increased energy in the alpha, beta, theta, and delta power bands in 12-month-old Q84 mice compared with wild-type mice^36^.

Given the sleep disorders reported in SCA3 patients and the above findings, we hypothesized that Q84 mice will show disrupted sleep structure and altered EEG spectral signatures during REM and NREM. Using 12-hour EEG recordings and spectrographic analysis, we sought to determine whether sleep is indeed disturbed in the Q84 mouse model of SCA3, and to evaluate whether parietal and frontal neural oscillations are altered in these mice in specific sleep states.

## METHODS

### Experimental Design

All animal procedures were approved by the University of Michigan Committee on the Use and Care of Animals (Protocols PRO00009818 and PRO00008409). Genotyping was performed by PCR on DNA isolated from tail biopsy at the time of weaning and post-mortem, as previously described^35^. We used Q84/Q84, Q84/WT, and WT/WT littermates. Mice of both sexes were used in all experiments: Q84/Q84 (N=11; 5 females, 6 males), Q84/WT (N=8; 1 female, 7 males), and WT/WT (N=12; 5 females, 7 males). As shown in our timeline (Figure 1A), first we weighed all mice, and assessed motor function to confirm the published phenotypic characteristics of Q84 mice^35^ and to investigate circadian differences among genotypes.

**Figure 1.**
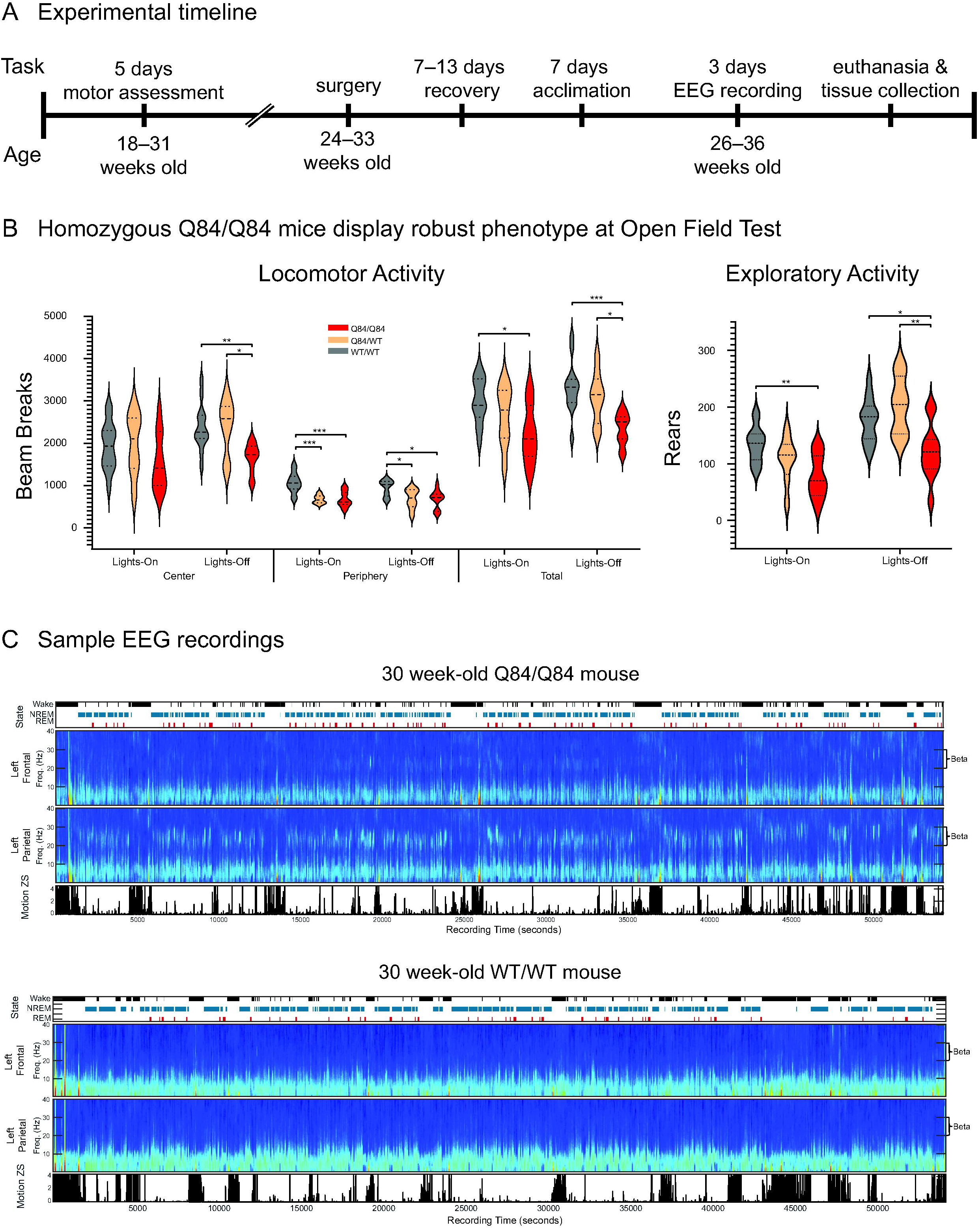
Measurement of motor behavior and sleep in SCA3 Q84 mice. **(A)** The motor behavior of 18–31-week-old mice was assessed in the morning with the lights on and in the evening with the lights off for 5 days. Next, mice underwent surgery to implant the recording electrodes at 24–33 weeks of age and, after 7–13 days of recovery, mice were acclimated to the recording equipment for one week. Finally, EEG recordings were obtained when the mice were 26–36-week-old for three consecutive days. **(B)** Homozygous Q84/Q84 mice (red, n=11) display decreased locomotor (beam breaks) and exploratory (number of rears) activities on the open-field test for 30 minutes compared with wildtype WT/WT mice (grey, n=13). Hemizygous Q84/WT mice (orange, n=8) display an intermediate phenotype. Statistical analysis conducted according to repeated measures ANOVA with Bonferroni-corrected post-hoc estimated marginal means test, * p<0.05, ** p <0.01, *** p<0.001. **(C)** Representative EEG recordings at the third day of evaluation of 30-week-old homozygous (top) and wild-type (bottom) mice. For each mouse there is a display of the hypnogram (black: wake, blue: non-rapid-eye movement sleep (NREM), red: rapid-eye movement sleep (REM)), spectrograms of the left frontal and parietal electrodes, and movement variance. Spectrograms show frequency power versus time from EEG recordings and this data was analyzed to determine differences between genotypes in oscillatory power during each of wake, REM and NREM. Increased beta power (20–30Hz) can be seen in the homozygous mouse during REM and NREM.

Following behavioral assessment, mice were split into groups of four for electrophysiology. Before surgery, mice were housed in cages with a maximum number of five animals, whereas post-surgery they were single housed. At all times mice were maintained in a standard 12-hour light/dark cycle (zeitgeber time (ZT) 0/12) with food and water *ad libitum*. One group had 12 weeks to adjust to a one-hour forward time shift due to daylight savings time. For each group, stereotaxic craniotomy surgeries were performed over one week (details below), which were followed by periods of recovery and acclimation to the recording equipment. Electrophysiologic recordings were then performed for three consecutive days while the mice behaved and slept freely. We analyzed data from the third recording day.

### Motor function evaluation

Motor function was assessed in 18–31-week-old mice, twice daily for five consecutive days using tests as previously described^35^. Each day, the first, lights-on, session started at approximately zeitgeber time (ZT) 3.5 and the second, lights-off, session started at ZT 15.5. In lights-off sessions, mice were only exposed to red wavelength lights. During the first four consecutive days (Days 1–4) we assessed performance on round and square balance beams followed by the rotarod test. On Day 5, we evaluated locomotor and exploratory activities in an open field chamber for 30 minutes.

For balance-beam testing, mice crossed a 44-cm-long plexiglass beam from an open clear platform to an enclosed black platform both placed at 53 cm of height^35^ for two consecutive trials on each of the four consecutive days and nights. Mice crossed first a 1 cm diameter round beam, and then a 0.5 cm-wide square beam. Time to traverse the beam was recorded for each trial with a 20-second maximum cutoff, and falls were scored as 20 seconds.

For rotarod testing, mice were placed on a rod accelerating from 4 to 40 revolutions per minute (RPM) over the course of 300 seconds (ENV-574M, Med Associates Inc.). Each animal performed two trials in the lights-on session and two trials in the lights-off session for four consecutive days. Mice rested for at least 30 minutes between the two trials. For each trial, the latency to fall off the rod was recorded, and was recorded as 300 seconds for animals that did not fall.

For the open field test^35^, each animal was placed once in the lights-on session and once in the lights-off session in the center of an open-field apparatus with XYZ infrared beams (San Diego Instruments). The number of beam brakes (locomotor activity) and rears (exploratory activity) was recorded for 30 minutes.

### Electrophysiology

Surgeries were performed on 24–33-week-old mice after testing for motor performance. We induced anesthesia with 4% isoflurane, administered carprofen IP for analgesia, and then placed the mouse in a stereotaxic frame, maintaining 0.5–2% isoflurane-induced anesthesia via nose cone. We also administered bupivacaine along the scalp midline before incision. After scalp incision, we drilled six holes, two for the frontal electrodes at +2 anterior/posterior (AP) and +/-1.25 medial/lateral (ML), two for the parietal electrodes at −2 AP and +/-1.25 ML, and two on the cerebellum, just below the lambda at +/-1.25 ML. In the holes, we placed 4–5 mm screws with already soldered wires. These wires were then soldered to a 32 or 64 channel Omnetics connector. The assembly was fixed to the screws and skull using acrylic dental cement, Unifast Trad. A copper mesh cage was then placed around the assembly. The cerebellar electrode at +1.25 ML was soldered to the copper cage to act as ground. After two Q84/Q84 mice died shortly after the surgery, we administered intraperitoneally methylprednisolone (30 mg/kg) to the rest of the mice to increase survival rate. Overall, we observed increased mortality in transgenic mice post-surgery that may relate to the previously observed genotype-based sensitivity^35^. After recovering for 7–13 days following surgery, mice were acclimated to their individual boxes in the recording room and were plugged in to the EEG recording equipment daily for one week, starting from 1 hour and gradually increasing to 10 hours.

We successfully obtained EEG recordings from 26–36-week-old Q84/Q84 (N=5), 3 Q84/WT (N=3), and WT/WT mice (N=9). EEG recordings were collected for three consecutive days in boxes grounded using aluminum foil. The first two days served as further acclimation to the recording procedure and thus were not analyzed. Daily, mice were transferred from the housing room to the recording room, and while the lights were still off, each mouse was placed into its recording box. Recordings started about 10 minutes before the lights were on and stopped about three hours after lights were off. Recordings were performed at 20,000Hz using an Intan RHD2000 Evaluation board and a preamplifier with an accelerometer chip to record movement in three dimensions. Mice were allowed to sleep and behave freely during these sessions. We recorded up to four mice at the same time.

### Sleep and EEG Analysis

Sleep scoring was performed by an automated MATLAB algorithm (SleepScoreMaster.m) and manually checked (TheStateEditor.m) from the ’buzcode’ library (https://github.com/buzsakilab/buzcode) as in previous work^37^. Two WT/WT mice had one electrode each (one left parietal and one left frontal) excluded from all analyses due to lack of signal. Spectrographic power, theta ratio and slow-wave activity, and movement estimates, based on the variance of the accelerometer sensors, were used to segregate REM, NREM, and wake brain states. Further analysis was done using custom code in MATLAB.

Here, we analyzed data from the third day of recordings, when the lights were on and mice typically spend more time sleeping. In addition to the total amount of time spent in each brain state, we also assessed sleep fragmentation by counting the number of bouts for each state and calculating their average duration. For spectral power analysis, we excluded extreme outliers separately for each channel by removing any seconds with elements that were more than 15 interquartile ranges above the upper quartile or below the lower quartile. Next, the power data for each frequency band of interest was normalized by dividing over the median value in that band over the 12-hour lights-on period. Before quantification, the power bands were defined as: delta 0.5–4Hz, theta 5–10Hz, spindle 11–19Hz, beta 20–30Hz, gamma 40–100Hz, and ripple 130–180Hz. Finally, to create average spectra we calculated the mean normalized spectral power per band for each brain state in the frontal and parietal electrodes.

### Statistics

Statistical analysis of the behavioral tests was performed on SPSS using repeated measures ANOVAs and *post-hoc* estimated marginal means, with day, time of day, and trial, where appropriate, as within-subjects factors and genotype as between-subjects factor. Kruskal Wallis tests and *post-hoc* pairwise comparisons were used to assess sleep stage metrics and spectral power differences between genotypes. Mann-Whitney tests were used to assess for differences between two genotypes. For all tests, the level of significance was determined as p<0.05. All *post-hoc* tests were Bonferroni corrected.

## RESULTS

### SCA3 mice show similar motor impairment in the lights-on phase to previous findings

We tested SCA3 Q84 mice at 18–31 weeks of age, when overt motor impairment has been previously reported^35^. First, we confirmed that Q84/Q84 mice replicated previous findings^35^ in loss of weight gain and motor impairments evaluated in the lights-on phase (Figure 1 and Supplementary Figure 1). We found a statistically significant main effect of the genotype in the round balance beam performance and all measures of open field behavior but not in the square balance beam, or the rotarod (Figure 1 and Supplementary Figure 1). Q84/Q84 mice were slower to traverse the round balance beam (Supplementary Figure 1A) and displayed fewer beam breaks and rears in the open field test (Figure 1B) than WT/WT mice. Q84/WT mice showed fewer beam breaks than WT/WT mice at the periphery of the open field (Figure 1B). Moreover, in some trials on the first two days, Q84/Q84 mice were slower to traverse the square balance beam and fell faster from the rotarod than WT/WT mice (Supplementary Figure 1B and C). Overall, while the 18–31-week-old homozygous Q84/Q84 mice used in this study showed robust loss of weight gain and motor dysfunction, the hemizygous Q84/WT mice revealed mild motor impairment compared with WT/WT littermates.

### SCA3 mice show a trend for circadian differences

Given the lack of previous circadian studies of motor behavior in SCA3 mouse models, here we investigated movement symptoms in SCA3 Q84 transgenic mice during both light and dark phases of the 24-hour cycle. In line with the nocturnal nature of lab mice, we found a main effect of the time of the day (lights-on or off) in all tasks with all the mice being more active during the lights-off sessions, except in the peripheral area of the open field apparatus (Figure 1B). Whereas we did not observe any overall statistically significant interactions between genotype and time of the day, *post-hoc* tests indicated some circadian differences for each genotype (Figure 1, Supplementary Figure 1): 1) in the round balance beam, only Q84/Q84 mice traversed the balance beam faster in the lights-off session (p=0.01); 2) in contrast, only the WT/WT mice stayed longer on the rotarod (p<0.001) and had more beam breaks in the central area of the open-field (p=0.025) during the lights-off session; and 3) all three genotypes demonstrated higher exploratory activity during the lights-off session (all p<0.01).

### SCA3 mice display increased REM duration and sleep/wake fragmentation compared to wild-type mice

After examining the circadian aspects of the motor phenotype, we studied the impacts of the *ATXN3* CAG expansion on sleep in Q84 mice by conducting EEG recordings in 26–36-week-old mice continuously for 12 hours during the lights-on period, when mice typically sleep. We used software with manual verification to score each one second period as either REM, NREM or wake. In terms of total duration of each state, we only observed an increase of total REM duration in Q84/Q84 mice compared with WT/WT mice (Figure 2). Importantly, we found that Q84/Q84 mice showed an increased number of bouts and shorter bout duration of REM, NREM, and WAKE compared to WT/WT mice, all p<0.05 (Figure 2), indicating increased sleep fragmentation in these mutant mice, similar to previous observations in SCA3 subjects^28, 29, 38^.

**Figure 2.**
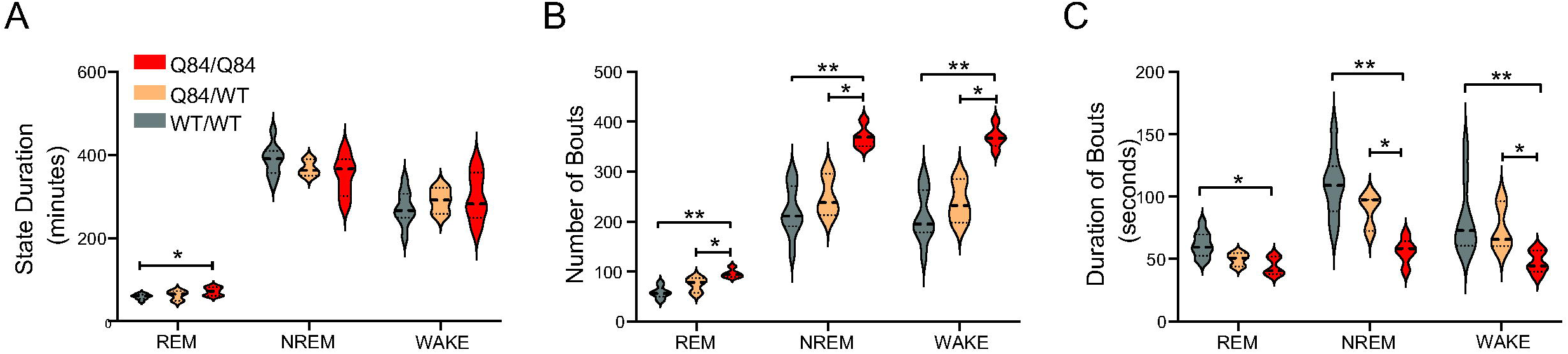
SCA3 Q84 mice show increased REM duration and sleep/wake fragmentation. **(A)** Homozygous Q84/Q84 mice (red, n=5) reveal longer REM duration overall than wild-type WT/WT (grey, n=9) littermates, according to unadjusted Mann Whitney. **(B)** Homozygous mice show more bouts of each state than their wild-type and hemizygous Q84/WT (orange, n=3) littermates. **(C)** In all states, homozygous mice display shorter bouts than wild-type mice. In NREM and wake, homozygous mice have shorter bouts than hemizygous mice. Thus, homozygous mice show that both sleep and wake states are composed of more bouts of shorter duration than wild-type mice. All violin plots display results from the 12 hours lights-on period (zeitgeber time (ZT) 0–12). Tick-down lines: statistical difference according to Kruskal Wallis with Bonferroni corrected post-hoc tests. Capped lines: statistical difference only according to unadjusted Mann Whitney tests. * p<0.05, ** p <0.01, REM: rapid-eye movement sleep, NREM: non-REM.

### SCA3 mice display altered neural oscillations during REM and NREM sleep

Many movement disorders including the synucleinopathies display altered oscillatory states during sleep relative to healthy subjects, even preceding clinical disease onset^39, 40^. To quantify differences in brain EEG oscillations between SCA3 mice and controls, we used spectrographic analysis to quantify the power of all EEG oscillations across sleep (REM and NREM) and wake states (Figure 3, 4, and 5). Visual examination of the spectrograms of all Q84/Q84 mice revealed the presence of beta power during REM and NREM (Figure 1C).

**Figure 3.**
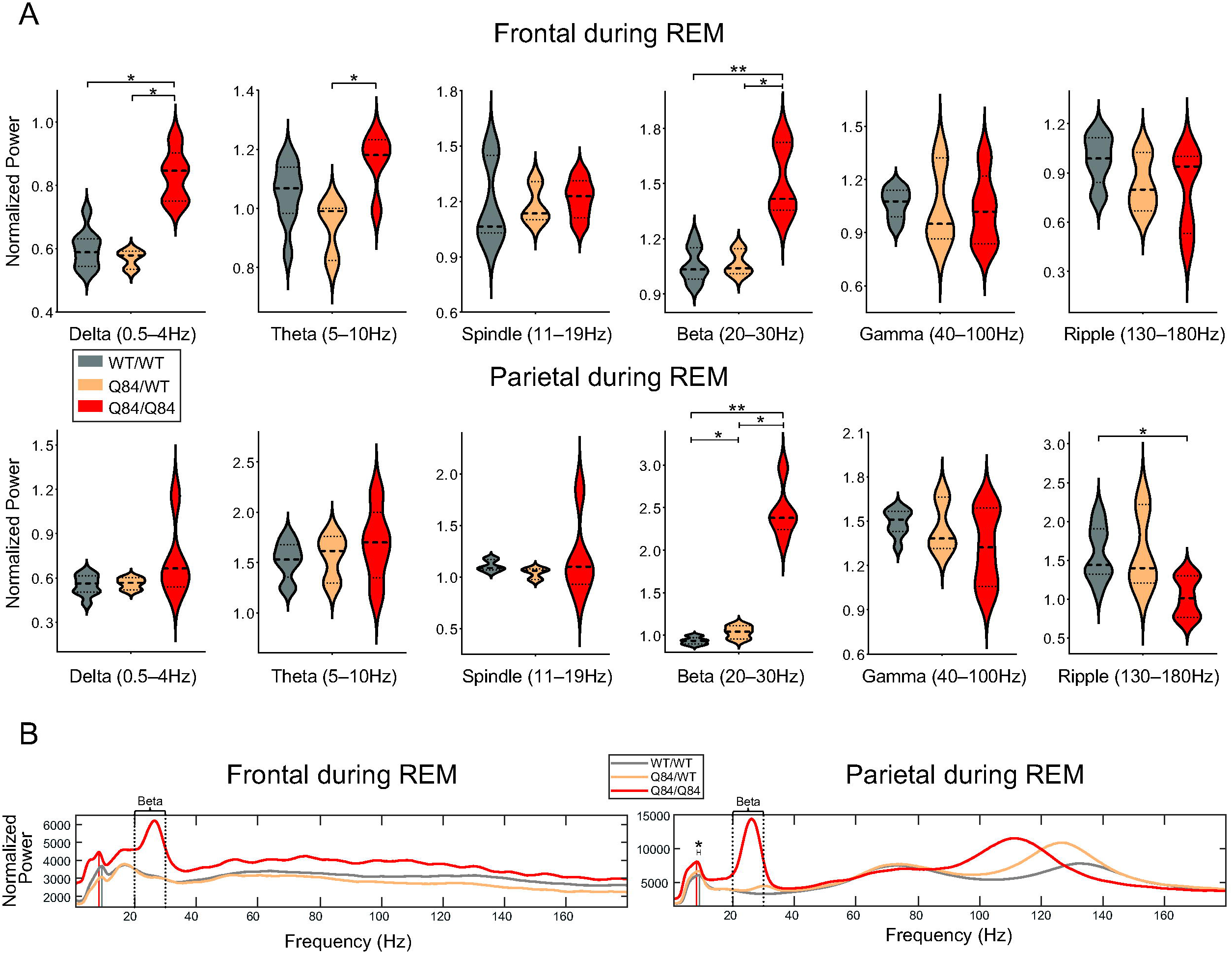
In REM sleep, homozygous Q84/Q84 mice show altered oscillatory power across many frequency bands, most clearly in frontal sites. **(A)** Homozygous Q84/Q84 mice (red, n=5) show higher oscillatory power compared to the wild-type WT/WT (grey, n=9) and/or hemizygous Q84/WT (orange, n=3) littermates in the delta, theta, and beta frequency bands in frontal sites during REM. In parietal sites during REM, homozygous mice show higher beta and lower ripple activity than the wild-type mice, and hemizygous mice show an intermediate phenotype in the beta band. **(B)** The total power averaged across all seconds of REM for each genotype plotted for better visualization. Homozygous mice show higher power across the low frequency bands including delta, theta, spindle, and beta bands spanning from 0.5–30Hz in both frontal and parietal electrodes. Vertical grey and red lines indicate the peak theta frequency for wild-type and homozygous mice. In the parietal sites, theta peak is slower in the homozygous mice than in the wild-type mice. Frontal sites show higher power in all frequencies in fact, but parietal sites do not show overall changes in gamma power, rather only a frequency shift of a power bump from around 135Hz to 110Hz. All plots display results from the 12 hours lights-on period (zeitgeber time (ZT) 0–12) and power is normalized by dividing over the respective frequency’s median value. Tick-down lines, statistical difference according to Kruskal Wallis with Bonferroni corrected post-hoc test. Capped lines, statistical difference only according to unadjusted Mann Whitney tests. * p<0.05, ** p <0.01. REM: rapid-eye movement sleep.

**Figure 4.**
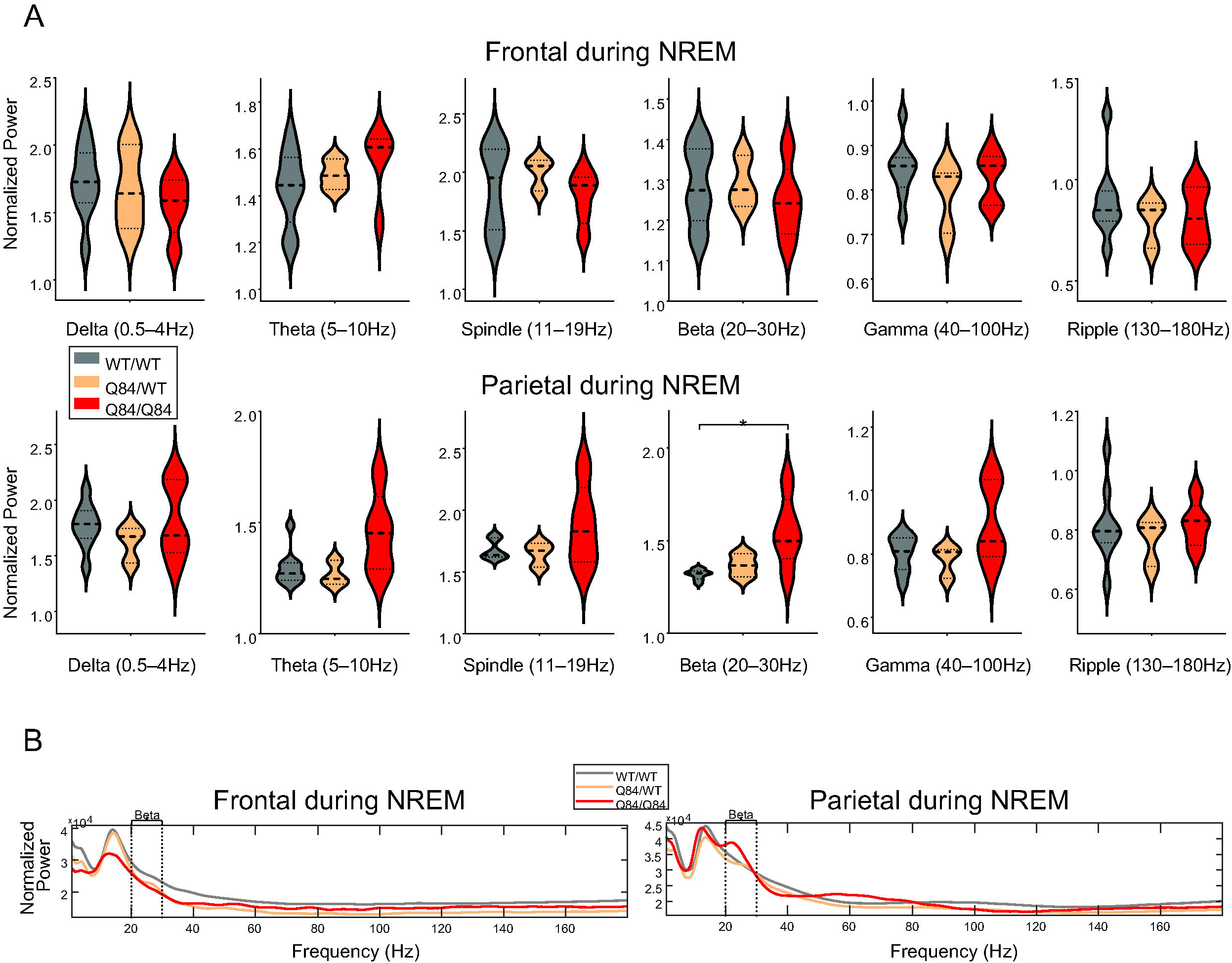
In NREM sleep, homozygous Q84/Q84 mice show increased parietal beta power. **(A)** Homozygous Q84/Q84 mice (red, n=5) show higher oscillatory power compared to the wild-type WT/WT (grey, n=9) in the parietal beta band power during NREM. **(B)** The total power averaged across all seconds of NREM for each genotype plotted for better visualization. Trends towards reduced delta and spindle power can be appreciated – both thalamocortical rhythms. All plots display results come from the 12 hours lights-on period (zeitgeber time (ZT) 0–12) and power is normalized by dividing over the respective frequency’s median value. Tick-down lines: statistical difference according to Kruskal Wallis with Bonferroni corrected *post-hoc* test. (red: homozygous Q84/Q84 mice (n=5), orange: hemizygous Q84/WT mice (n=3), grey: wild-type WT/WT mice (n=9)) NREM: non-rapid-eye movement sleep, * p<0.05

**Figure 5.**
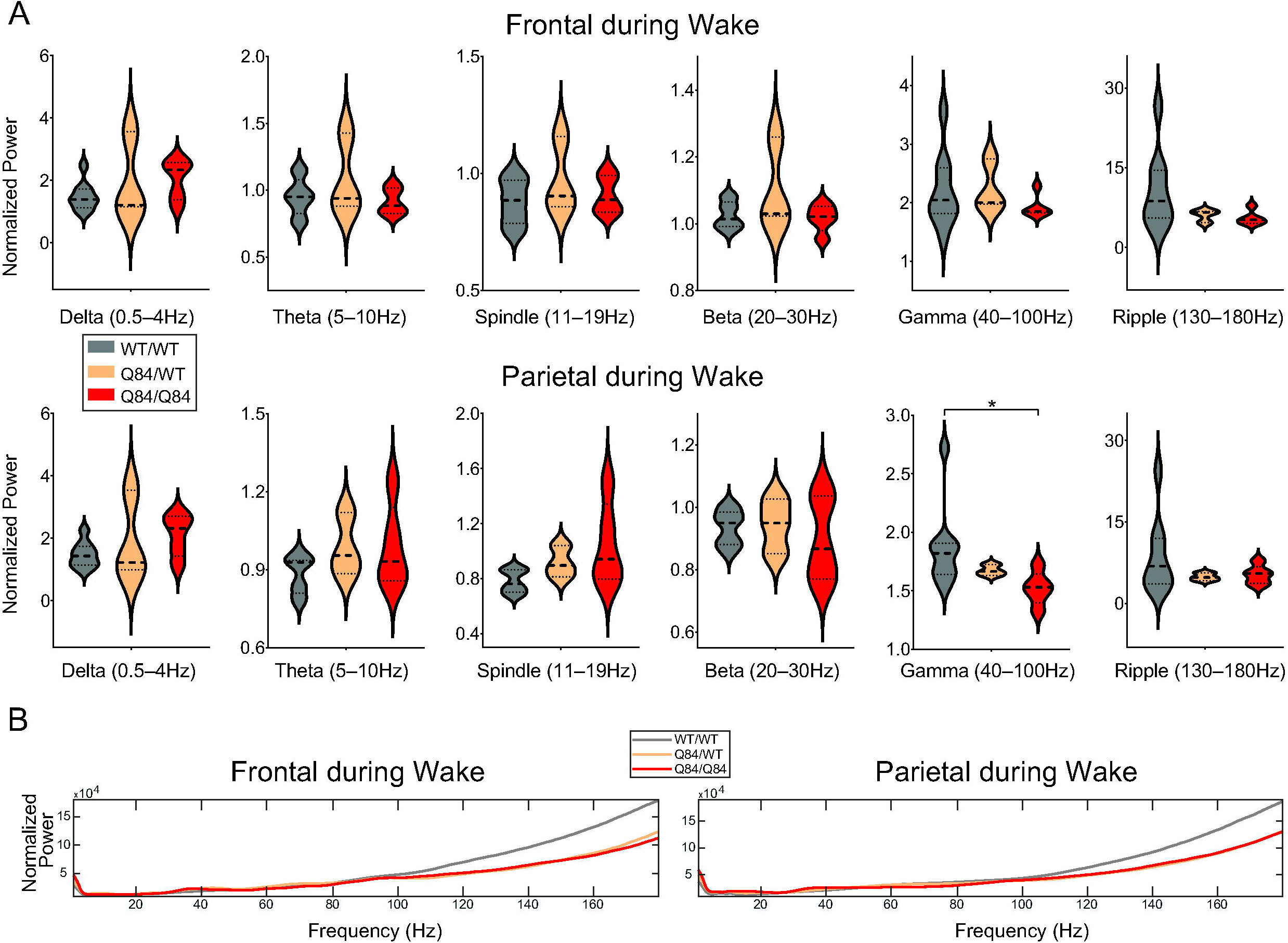
In wake, homozygous Q84/Q84 mice show decreased parietal gamma power. **(A)** Homozygous Q84/Q84 mice (red, n=5) show decreased gamma power in the parietal area during wake when compared to their wild-type WT/WT (grey, n=9) littermates during wake. **(B)** The total power averaged across all seconds of wake for each genotype plotted for better visualization. Normalization by dividing over the total median value across the full recording was kept to allow for consistency with data in part A, but that creates the upward trend towards higher frequency bands. All plots display results that come from the 12 hours lights-on period (zeitgeber time (ZT) 0–12) and power is normalized by dividing over the respective frequency’s median value. Tick-down lines, statistical difference according to Bonferroni corrected *post-hoc* Kruskal Wallis tests. Significance as * p<0.05.

First, we validated our visual finding of increased beta power in Q84/Q84 mice compared to WT/WT mice during REM and NREM (Figure 3 and Figure 4): whereas during REM, beta power was increased in both frontal and parietal regions (Figure 3), during NREM it was only increased in the parietal area (with significance set at p<0.05 by Kruskal Wallis) (Figure 4). We also observed an intermediate phenotype in Q84/WT mice during REM (Figure 3): in the parietal electrodes, Q84/WT showed higher beta power compared to WT/WT mice and lower compared to Q84/Q84 mice (Figure 3); and in the frontal electrodes, Q84/WT mice displayed lower beta power than Q84/Q84 mice (Figure 3).

We verified additional cross-genotype anatomy-specific alterations of brain oscillations during REM. During REM, compared with WT/WT mice theQ84/Q84 mice revealed higher frontal delta power (Figure 3), lower parietal ripple power (Figure 3), higher frontal theta band power (Figure 3), and slower peak frequency of theta rhythm in the parietal area (Figure 3). Moreover, during REM Q84/Q84 mice showed increased power in the frontal delta and theta bands compared to Q84/WT mice (Figure 3). Finally, during wake, we observed lower parietal gamma oscillation in Q84/Q84 mice compared to WT/WT littermates (Figure 5).

## DISCUSSION

In this study we sought to identify alterations in sleep architecture and EEG spectral power in motor impaired SCA3 Q84 transgenic mice that are frequently used in preclinical trials for SCA3^35^. EEG analysis of 26–36-week-old homozygous and hemizygous Q84 mice and wild-type littermates allowed us to identify quantitative sleep changes in these mice: a) increased fragmentation in all states and REM duration; b) higher beta power during REM and NREM; and c) several alterations of brain EEG oscillations during REM and wake including in the theta and delta bands. Importantly, both Q84 mice and patients with SCA3 have alterations in REM and NREM sleep.

As expected, 18–31-week-old Q84/Q84 mice displayed motor impairments and reduced weight gain. The novel circadian differences in some genotypes in our work may be partially due to the difficulty of each task in combination with the degree of motor impairment, and/or due to learning defects.

In Q84/Q84 mice, we found that sleep architecture displayed increased fragmentation in REM, NREM, and wake periods. Importantly, this mirrors findings of sleep fragmentation^15, 29, 41^ and decreased sleep efficiency^29^ previously identified in SCA3 patients. Such sleep fragmentation can disrupt the homeostatic functions of sleep and thus have a direct impact on proper neurobiological function, including impairment of protein homeostasis and decreased clearance of toxic proteins^42, 43^. The detrimental effects of chronic sleep fragmentation were observed in an AD mouse model where sleep fragmentation increased amyloid β deposition^44^. The sleep fragmentation we observed seems to be a common feature in neurodegenerative diseases^45–52^ and could, therefore, contribute to improper clearance of the misfolded and aggregation-prone mutant ATXN3 protein in SCA3.

We found increased REM duration in Q84/Q84 mice. In SCA3 patients, REM findings are mixed. One study (n=15) showed decreased REM duration^29^, but a larger study (n=47) showed no changes in REM duration^28^. Patients in both studies had similar age at the evaluation time, age at disease onset, and size of the expanded CAG repeat but different number of patients and sex ratios. Overall, REM appears to be affected by SCA3. How REM is dysregulated may depend on which areas of the brainstem and cerebellum are affected by SCA3 neuronal loss^5, 6^ since they both contain circuits controlling REM^53–57^. Age may also play a role in this progressive disease, and longitudinal imaging and sleep studies assessing REM at different disease stages in individuals with SCA3 and SCA3 animal models are needed.

While beta-band changes were most prominent during REM, we found additional EEG alterations of delta and theta band powers and theta band frequency. These results are similar to previous EEG findings in 12-month-old SCA3 Q84 mice^36^, although it is unclear in which alertness state these findings occurred. We also observed a decrease in ripple power during REM and an increase of gamma power during wake in homozygous mice. While there have been findings of decreased spindle (11–16Hz) density in SCA3 patients^33^, in our 6.5–9-month-old SCA3 Q84 mouse model we did not observe any significant differences in spindle power (11–19Hz).

Most prominently, we showed increased beta power during NREM and REM sleep in Q84 mice. The degrees and significances of this change usually correlated with the dose of the *ATXN3* transgene in the Q84 mice. In Q84/Q84 mice, higher beta power was seen across all anatomical locations during REM but only in the parietal electrodes during NREM. In Q84/WT mice, of which we only had 3, an intermediate phenotype was seen during REM in the parietal electrodes. Increased beta in the motor cortex has been reported before in this SCA3 mouse model^36^, although it is unclear in which sleep state. Based on mouse brain anatomy^58, 59^, the parietal electrodes are closest to the basal ganglia and thus more likely to capture signals from that area than the frontal electrodes. This is significant since the basal ganglia are known to be affected by neuronal loss and ATXN3 aggregation in SCA3 patients^6^. The increased beta power during REM and NREM in Q84 mice is reminiscent of increased beta oscillations observed in other neurodegenerative diseases and sleep disorders. Alterations, most frequently increases, in beta oscillations have been observed during wake and sleep states in patients or animal models of PD^60–68^, HD^69, 70, 71–78, 79–84^, dementia with Lewy bodies^85, 86^, insomnia^60, 87^, and RBD^63, 88–90^. In PD, beta oscillations in the basal ganglia are usually pathological^60, 62, 64, 67, 68, 91^ and appear to be linked to basal ganglia dysfunction or compensation. Thus, the triad of RBD, beta oscillations, and basal ganglia appears to be a unifying feature of these diseases, including SCA3.

Clinically, RBD is considered a predictor of some neurodegenerative disorders, including synucleinopathies and dementias^92, 93^. It is known that SCA3 patients experience RBD and other sleep changes^94^. Moreover, beta event-related synchronization at a motor task appears to be reduced in SCA3 patients in comparison to healthy controls^95^, but it is yet to be determined whether SCA3 patients also show increased beta activity during REM. These findings indicate that the combination of RBD and basal ganglia-based beta-oscillatory changes could represent an overlap in meso-scale pathology across SCA3 and other neurodegenerative disorders. This change may represent a primary alteration or compensation for other pathology.

In summary, we found that sleep architecture and specific brain oscillations, in particular beta power, are altered in motor impaired SCA3 Q84 mice. The sleep EEG abnormalities identified here may point to altered neuronal circuits pertinent to SCA3, such as the basal ganglia, yielding greater mechanistic detail of SCA3 pathogenesis, and should prompt additional human studies of spectral changes both in and out of sleep. Many of our EEG spectral findings were both state and anatomy-specific; accordingly, future deep-tissue recordings would be needed to better understand the anatomical and spiking implications of the observed regional alterations in SCA3 mice. Limitations here include EEG recordings from few hemizygous mice and possible discrepancies in REM duration findings in mice versus human, though human results are mixed. Also, longitudinal sleep EEG studies throughout the lifespan of people with SCA3 and transgenic mouse models will be needed to extend our observations on sleep architecture and spectral alterations, determine their relation to progression of motor symptoms and brain pathophysiology, and evaluate their potential translational utility. Hence, specific quantifiable sleep and EEG changes may serve as a target for the development of disease-modifying therapies and as disease biomarkers in SCA3.

## ACKNOWLEDGEMENTS

The authors would like to thank: Louise O’Brien, Lezio Soares Bueno, and Paul Fitzgerald for illuminating discussions and training.

## AUTHOR CONTRIBUTIONS

MET, MCC, and BOW designed and supervised the experimental work; MCC and BOW provided the funding for the experiments; MET, AG, AJB, and RW performed the experiments; MET, MCC, and BOW analyzed the results; MET, MCC, BOW, and HLP wrote the manuscript; all authors participated in the discussion and revised the manuscript.

## FINANCIAL DISCLOSURE (for the preceding 12 months)

The authors have no financial disclosures to declare for the past 12 months.

## REFERENCES

1. Ruano, L., Melo, C., Silva, M. C. & Coutinho, P. The global epidemiology of hereditary ataxia and spastic paraplegia: A systematic review of prevalence studies. Neuroepidemiology vol. 42 174–183 (2014).

2. Klockgether, T., Mariotti, C. & Paulson, H. L. Spinocerebellar ataxia. Nat. Rev. Dis. Prim. 2019 51 **5**, 1–21 (2019).

3. Bettencourt, C., Santos, C., Montiel, R., et al. Increased transcript diversity: Novel splicing variants of Machado-Joseph Disease gene (ATXN3). Neurogenetics 11, 193–202 (2010).

4. ATXN3 ataxin 3 [Homo sapiens (human)] - Gene - NCBI. https://www.ncbi.nlm.nih.gov/gene/4287.

5. Rüb, U., Schöls, L., Paulson, H., et al. Clinical features, neurogenetics and neuropathology of the polyglutamine spinocerebellar ataxias type 1, 2, 3, 6 and 7. Progress in Neurobiology vol. 104 38–66 (2013).

6. Seidel, K., Siswanto, S., Brunt, E. R. P., Den Dunnen, W., Korf, H. W. & Rüb, U. Brain pathology of spinocerebellar ataxias. Acta Neuropathol. 2012 1241 124, 1–21 (2012).

7. McLoughlin, H. S., Moore, L. R. & Paulson, H. L. Pathogenesis of SCA3 and implications for other polyglutamine diseases. Neurobiology of Disease vol. 134 (2020).

8. Costa, M. do C. & Paulson, H. L. Toward understanding Machado-Joseph Disease. Prog. Neurobiol. 97, 239 (2012).

9. Paulson, H. & Shakkottai, V. Spinocerebellar Ataxia Type 3. GeneReviews® (2020).

10. Pedroso, J. L., Braga-Neto, P., Martinez, A. R. M., et al. Sleep disorders in Machado-Joseph disease. Curr. Opin. Psychiatry 29, 402–408 (2016).

11. D’Abreu, A., Friedman, J. & Coskun, J. Non-movement disorder heralds symptoms of Macho-Joseph disease year before ataxia. Mov. Disord. 20, 739–741 (2005).

12. Fukutake, T., Shinotoh, H., Nishino, H., et al. Homozygous Machado-Joseph disease presenting as REM sleep behaviour disorder and prominent psychiatric symptoms. Eur. J. Neurol. 9, 97–100 (2002).

13. D’Abreu, A., França, M., Conz, L., et al. Sleep symptoms and their clinical correlates in Machado-Joseph disease. Acta Neurol. Scand. 119, 277–280 (2009).

14. Takei, A., Fukazawa, T., Hamada, T., et al. Effects of Tandospirone on ‘5-HT1A Receptor-Associated Symptoms’ in Patients with Machado-Josephe Disease An Open-Label Study. Clin. Neuropharmacol. 27, 9–13 (2004).

15. Kushida, C. A., Clerk, A. A., Kirsch, C. M., Hotson, J. R. & Guilleminault, C. Prolonged confusion with nocturnal wandering arising from NREM and REM sleep: A case report. Sleep 18, 757–764 (1995).

16. Friedman, J. H., Fernandez, H. H. & Sudarsky, L. R. REM behavior disorder and excessive daytime somnolence in Machado-Joseph disease (SCA-3). Mov. Disord. 18, 1520–1522 (2003).

17. Pedroso, J. L., Braga-Neto, P., Felício, A. C., et al. Sleep disorders in Machado-Joseph disease: Frequency, discriminative thresholds, predictive values, and correlation with ataxia-related motor and non-motor features. Cerebellum 10, 291–295 (2011).

18. Folha Santos, F. A., de Carvalho, L. B. C., Prado, L. F. do, do Prado, G. F., Barsottini, O. G. & Pedroso, J. L. Sleep apnea in Machado-Joseph disease: a clinical and polysomnographic evaluation. Sleep Med. 48, 23–26 (2018).

19. Martinez, A. R. M., Nunes, M. B., Faber, I., D’Abreu, A., Lopes-Cendes, Í. & França, M. C. Fatigue and Its Associated Factors in Spinocerebellar Ataxia Type 3/Machado-Joseph Disease. Cerebellum 16, 118–121 (2017).

20. Yang, J. S., Xu, H. L., Chen, P. P., et al. Ataxic Severity Is Positively Correlated With Fatigue in Spinocerebellar Ataxia Type 3 Patients. Front. Neurol. 11, (2020).

21. Abele, M., Bürk, K., Laccone, F., Dichgans, J. & Klockgether, T. Restless legs syndrome in spinocerebellar ataxia types 1, 2, and 3. J. Neurol. 248, 311–314 (2001).

22. Gitaí, L. L. G., Éckeli, A. L., Sobreira-Neto, M. A., et al. Which Factors in Spinocerebellar Ataxia Type 3 Patients Are Associated with Restless Legs Syndrome/Willis-Ekbom Disease? Cerebellum 20, 21–30 (2021).

23. Pedroso, J. L., Braga-Neto, P., Felício, A. C., et al. Sleep disorders in Machado-Joseph disease: A dopamine transporter imaging study. J. Neurol. Sci. 324, 90–93 (2013).

24. Reimold, M., Globas, C., Gleichmann, M., et al. Spinocerebellar ataxia type 1,2, and 3 and restless legs syndrome: Striatal dopamine D2 receptor status investigated by [11C] Raclopride positron emission tomography. Mov. Disord. 21, 1667–1673 (2006).

25. Schöls, L., Haan, J., Riess, O., Amoiridis, G. & Przuntek, H. Sleep disturbance in spinocerebellar ataxias: Is the SCA3 mutation a cause of restless legs syndrome? Neurology 51, 1603–1607 (1998).

26. Pedroso, J. L., Bezerra, M. L. E., Braga-Neto, P., et al. Is Neuropathy Involved with Restless Legs Syndrome in Machado-Joseph Disease? Eur. Neurol. 66, 200–203 (2011).

27. Pedroso, J. L., Bor-Seng-Shu, E., Braga-Neto, P., et al. Neurophysiological studies and non-motor symptoms prior to ataxia in a patient with Machado-Joseph disease: Trying to understand the natural history of brain degeneration. Cerebellum 13, 447–451 (2014).

28. Silva, G. M. F., Pedroso, J. L., Dos Santos, D. F., et al. NREM-related parasomnias in Machado-Joseph disease: Clinical and polysomnographic evaluation. J. Sleep Res. 25, 11–15 (2016).

29. Chi, N. F., Shiao, G. M., Ku, H. L. & Soong, B. W. Sleep disruption in spinocerebellar ataxia type 3: A genetic and polysomnographic study. J. Chinese Med. Assoc. 76, 25–30 (2013).

30. Friedman, J. H. Presumed rapid eye movement behavior disorder in Machado-Joseph disease (Spinocerebellar ataxia type 3). Mov. Disord. 17, 1350–1353 (2002).

31. Syed, B. H., Rye, D. B. & Singh, G. REM sleep behavior disorder and SCA-3 (Machado-Joseph disease). Neurology 60, 148 (2003).

32. Iranzo, A., Muñoz, E., Santamaria, J., Vilaseca, I., Milà, M. & Tolosa, E. REM sleep behavior disorder and vocal cord paralysis in Machado-Joseph disease. Mov. Disord. 18, 1179–1183 (2003).

33. Seshagiri, D. V., Botta, R., Sasidharan, A., et al. Assessment of Sleep Spindle Density among Genetically Positive Spinocerebellar Ataxias Types 1, 2, and 3 Patients. Ann. Neurosci. 25, 106–111 (2018).

34. Cemal, C. K., Carroll, C. J., Lawrence, L., et al. YAC transgenic mice carrying pathological alleles of the MJD1 locus exhibit a mild and slowly progressive cerebellar deficit. Hum. Mol. Genet. 11, 1075–1094 (2002).

35. Do Carmo Costa, M., Luna-Cancalon, K., Fischer, S., et al. Toward RNAi therapy for the polyglutamine disease Machado-Joseph disease. Mol. Ther. 21, 1898–1908 (2013).

36. Yu, Y., He, X., Zhao, Z., et al. Nonlinear analysis of local field potentials and motor cortex EEG in spinocerebellar ataxia 3. J. Clin. Neurosci. 59, 298–304 (2019).

37. Watson, B. O., Levenstein, D., Greene, J. P., Gelinas, J. N. & Buzsáki, G. Network Homeostasis and State Dynamics of Neocortical Sleep. Neuron 90, 839–852 (2016).

38. Huebra, L., Coelho, F. M., Filho, F. M. R., Barsottini, O. G. & Pedroso, J. L. Sleep Disorders in Hereditary Ataxias. Curr. Neurol. Neurosci. Rep. 19, (2019).

39. St Louis, E. K., Boeve, A. R. & Boeve, B. F. REM Sleep Behavior Disorder in Parkinson’s Disease and Other Synucleinopathies. Mov. Disord. 32, 645–658 (2017).

40. Bailey, G. A., Hubbard, E. K., Fasano, A., et al. Sleep disturbance in movement disorders: insights, treatments and challenges. J. Neurol. Neurosurg. Psychiatry 92, 723– 736 (2021).

41. dos Santos, D. F., Pedroso, J. L., Braga-Neto, P., et al. Excessive fragmentary myoclonus in Machado-Joseph disease. Sleep Med. 15, 355–358 (2014).

42. Tononi, G. & Cirelli, C. Sleep function and synaptic homeostasis. Sleep Med. Rev. 10, 49–62 (2006).

43. Frank, M. G. & Heller, H. C. The Function(s) of Sleep. Handb. Exp. Pharmacol. 253, 3–34 (2019).

44. Minakawa, E. N., Miyazaki, K., Maruo, K., et al. Chronic sleep fragmentation exacerbates amyloid β deposition in Alzheimer’s disease model mice. Neurosci. Lett. 653, 362–369 (2017).

45. Cai, G. E., Luo, S., Chen, L. N., Lu, J. P., Huang, Y. J. & Ye, Q. Y. Sleep fragmentation as an important clinical characteristic of sleep disorders in Parkinson’s disease: a preliminary study. Chin. Med. J. (Engl). 132, 1788–1795 (2019).

46. Lim, A. S. P., Fleischman, D. A., Dawe, R. J., et al. Regional Neocortical Gray Matter Structure and Sleep Fragmentation in Older Adults. Sleep 39, 227–235 (2016).

47. Wennberg, A. M. V., Wu, M. N., Rosenberg, P. B. & Spira, A. P. Sleep Disturbance, Cognitive Decline, and Dementia: A Review. Semin. Neurol. 37, 395–406 (2017).

48. Xie, Y., Ba, L., Wang, M., et al. Chronic sleep fragmentation shares similar pathogenesis with neurodegenerative diseases: Endosome-autophagosome-lysosome pathway dysfunction and microglia-mediated neuroinflammation. CNS Neurosci. Ther. 26, 215– 227 (2020).

49. Owen, J. E. & Veasey, S. C. Impact of sleep disturbances on neurodegeneration: Insight from studies in animal models. Neurobiol. Dis. 139, (2020).

50. Dufort-Gervais, J., Mongrain, V. & Brouillette, J. Bidirectional relationships between sleep and amyloid-beta in the hippocampus. Neurobiol. Learn. Mem. 160, 108–117 (2019).

51. Fernandes, M., Chiaravalloti, A., Manfredi, N., et al. Nocturnal Hypoxia and Sleep Fragmentation May Drive Neurodegenerative Processes: The Compared Effects of Obstructive Sleep Apnea Syndrome and Periodic Limb Movement Disorder on Alzheimer’s Disease Biomarkers. J. Alzheimers. Dis. 1–13 (2022) doi:10.3233/JAD-215734.

52. Abbott, S. M. & Videnovic, A. Chronic sleep disturbance and neural injury: links to neurodegenerative disease. Nat. Sci. Sleep 8, 55–61 (2016).

53. Cunchillos, J. D. & De Andrés, I. Participation of the cerebellum in the regulation of the sleep-wakefulness cycle. Results in cerebellectomized cats. Electroencephalogr. Clin. Neurophysiol. 53, 549–558 (1982).

54. DelRosso, L. M. & Hoque, R. The cerebellum and sleep. Neurol. Clin. 32, 893–900 (2014).

55. Canto, C. B., Onuki, Y., Bruinsma, B., van der Werf, Y. D. & De Zeeuw, C. I. The Sleeping Cerebellum. Trends Neurosci. 40, 309–323 (2017).

56. Luppi, P. H., Clement, O., Sapin, E., et al. Brainstem mechanisms of paradoxical (REM) sleep generation. Pflugers Arch. 463, 43–52 (2012).

57. Fraigne, J. J., Torontali, Z. A., Snow, M. B. & Peever, J. H. REM Sleep at its Core - Circuits, Neurotransmitters, and Pathophysiology. Front. Neurol. 6, (2015).

58. Paxinos, G. & Franklin, K. B. J. The Mouse Brain in Stereotaxic Coordinates. Academic Press vol. 3rd Editio (2008).

59. Wang, Q., Ding, S. L., Li, Y., et al. The Allen Mouse Brain Common Coordinate Framework: A 3D Reference Atlas. Cell 181, 936–953.e20 (2020).

60. Mizrahi-Kliger, A. D., Kaplan, A., Israel, Z., Deffains, M. & Bergman, H. Basal ganglia beta oscillations during sleep underlie Parkinsonian insomnia. Proc. Natl. Acad. Sci. U. S. A. 117, 17359–17368 (2020).

61. Urrestarazu, E., Iriarte, J., Alegre, M., et al. Beta activity in the subthalamic nucleus during sleep in patients with Parkinson’s disease. Mov. Disord. 24, 254–260 (2009).

62. Baumgartner, A. J., Kushida, C. A., Summers, M. O., Kern, D. S., Abosch, A. & Thompson, J. A. Basal Ganglia Local Field Potentials as a Potential Biomarker for Sleep Disturbance in Parkinson’s Disease. Front. Neurol. 12, (2021).

63. Hackius, M., Werth, E., Sürücü, O., Baumann, C. R. & Imbach, L. L. Electrophysiological evidence for alternative motor networks in REM sleep behavior disorder. J. Neurosci. 36, 11795–11800 (2016).

64. Yu, Y., Sanabria, D. E., Wang, J., et al. Parkinsonism Alters Beta Burst Dynamics across the Basal Ganglia-Motor Cortical Network. J. Neurosci. 41, 2274–2286 (2021).

65. Gong, R., Wegscheider, M., Mühlberg, C., et al. Spatiotemporal features of β-γ phase-amplitude coupling in Parkinson’s disease derived from scalp EEG. Brain 144, 487–503 (2021).

66. Ghilardi, M. F., Tatti, E. & Quartarone, A. Beta power and movement-related beta modulation as hallmarks of energy for plasticity induction: Implications for Parkinson’s disease. Parkinsonism Relat. Disord. 88, 136–139 (2021).

67. Eisinger, R. S., Cagle, J. N., Opri, E., et al. Parkinsonian Beta Dynamics during Rest and Movement in the Dorsal Pallidum and Subthalamic Nucleus. J. Neurosci. 40, 2859 (2020).

68. Devergnas, A., Pittard, D., Bliwise, D. & Wichmann, T. Relationship between oscillatory activity in the cortico-basal ganglia network and parkinsonism in MPTP-treated monkeys. Neurobiol. Dis. 68, 156–166 (2014).

69. Morton, A. J. Circadian and sleep disorder in Huntington’s disease. Exp. Neurol. 243, 34– 44 (2013).

70. Fisher, S. P., Black, S. W., Schwartz, M. D., et al. Longitudinal analysis of the electroencephalogram and sleep phenotype in the R6/2 mouse model of Huntington’s disease. Brain 136, 2159–2172 (2013).

71. Kantor, S., Szabo, L., Varga, J., Cuesta, M. & Morton, A. J. Progressive sleep and electroencephalogram changes in mice carrying the Huntington’s disease mutation. Brain 136, 2147–2158 (2013).

72. Jeantet, Y., Cayzac, S. & Cho, Y. H. β oscillation during slow wave sleep and rapid eye movement sleep in the electroencephalogram of a transgenic mouse model of Huntington’s disease. PLoS One 8, (2013).

73. Neutel, D., Tchikviladzé, M., Charles, P., et al. Nocturnal agitation in Huntington disease is caused by arousal-related abnormal movements rather than by rapid eye movement sleep behavior disorder. Sleep Med. 16, 754–759 (2015).

74. Piano, C., Losurdo, A., Marca, G. Della, et al. Polysomnographic findings and clinical correlates in Huntington disease: A cross-sectional cohort study. Sleep 38, 1489–1495 (2015).

75. Lazar, A. S., Panin, F., Goodman, A. O. G., et al. Sleep deficits but no metabolic deficits in premanifest Huntington’s disease. Ann. Neurol. 78, 630–648 (2015).

76. Lebreton, F., Cayzac, S., Pietropaolo, S., Jeantet, Y. & Cho, Y. H. Sleep physiology alterations precede plethoric phenotypic changes in R6/1 Huntington’s disease mice. PLoS One 10, (2015).

77. Kantor, S., Varga, J. & Morton, A. J. A single dose of hypnotic corrects sleep and EEG abnormalities in symptomatic Huntington’s disease mice. Neuropharmacology 105, 298– 307 (2016).

78. Fisher, S. P., Schwartz, M. D., Wurts-Black, S., et al. Quantitative electroencephalographic analysis provides an early-stage indicator of disease onset and progression in the zQ175 knock-in mouse model of huntington’s disease. Sleep 39, 379– 391 (2016).

79. Piano, C., Imperatori, C., Losurdo, A., Bentivoglio, A. R., Cortelli, P. & Della Marca, G. Sleep-related modifications of EEG connectivity in the sensory-motor networks in Huntington Disease: An eLORETA study and review of the literature. Clin. Neurophysiol. 128, 1354–1363 (2017).

80. Piano, C., Mazzucchi, E., Bentivoglio, A. R., et al. Wake and Sleep EEG in Patients With Huntington Disease: An eLORETA Study and Review of the Literature. Clin. EEG Neurosci. 48, 60–71 (2017).

81. Piano, C., Marca, G. Della, Losurdo, A., et al. Subjective Assessment of Sleep in Huntington Disease: Reliability of Sleep Questionnaires Compared to Polysomnography. Neurodegener. Dis. 17, 330–337 (2017).

82. Zhang, Y., Ren, R., Yang, L., et al. Sleep in Huntington’s disease: a systematic review and meta-analysis of polysomongraphic findings. Sleep 42, (2019).

83. Smarr, B., Cutler, T., Loh, D. H., et al. Circadian dysfunction in the Q175 model of Huntington’s disease: Network analysis. J. Neurosci. Res. 97, 1606–1623 (2019).

84. Herzog-Krzywoszanska, R. & Krzywoszanski, L. Sleep disorders in Huntington’s disease. Front. Psychiatry 10, (2019).

85. Stylianou, M., Zaaimi, B., Thomas, A., Taylor, J. P. & LeBeau, F. E. N. Early Disruption of Cortical Sleep-Related Oscillations in a Mouse Model of Dementia With Lewy Bodies (DLB) Expressing Human Mutant (A30P) Alpha-Synuclein. Front. Neurosci. 14, (2020).

86. Morris, M., Sanchez, P. E., Verret, L., et al. Network dysfunction in α-synuclein transgenic mice and human Lewy body dementia. Ann. Clin. Transl. Neurol. 2, 1012–1028 (2015).

87. Spiegelhalder, K., Regen, W., Feige, B., et al. Increased EEG sigma and beta power during NREM sleep in primary insomnia. Biol. Psychol. 91, 329–333 (2012).

88. Figorilli, M., Lanza, G., Congiu, P., et al. Neurophysiological Aspects of REM Sleep Behavior Disorder (RBD): A Narrative Review. Brain Sci. 11, (2021).

89. Valomon, A., Riedner, B. A., Jones, S. G., et al. A high-density electroencephalography study reveals abnormal sleep homeostasis in patients with rapid eye movement sleep behavior disorder. Sci. Rep. 11, (2021).

90. Ferri, R., Rundo, F., Silvani, A., et al. REM Sleep EEG Instability in REM Sleep Behavior Disorder and Clonazepam Effects. Sleep 40, (2017).

91. Deffains, M. & Bergman, H. Parkinsonism-related β oscillations in the primate basal ganglia networks – Recent advances and clinical implications. Parkinsonism Relat. Disord. 59, 2–8 (2019).

92. Petit, D., Gagnon, J. F., Fantini, M. L., Ferini-Strambi, L. & Montplaisir, J. Sleep and quantitative EEG in neurodegenerative disorders. Journal of Psychosomatic Research vol. 56 487–496 (2004).

93. Zhang, F., Niu, L., Liu, X., et al. Rapid Eye Movement Sleep Behavior Disorder and Neurodegenerative Diseases: An Update. Aging Dis. 11, 315 (2020).

94. Moro, A., Moscovich, M., Farah, M., Camargo, C. H. F., Teive, H. A. G. & Munhoz, R. P. Nonmotor symptoms in spinocerebellar ataxias (SCAs). Cerebellum and Ataxias 6, 12 (2019).

95. Aoh, Y., Hsiao, H. J., Lu, M. K., et al. Event-Related Desynchronization/Synchronization in Spinocerebellar Ataxia Type 3. Front. Neurol. 10, (2019).

