## Supplemental Figure 1 for "Sleep alterations in a mouse model of Spinocerebellar ataxia type 3"

**SUPPLEMENTARY MATERIAL**

**Supplementary Figure 1. Motor impairment and decreased weight in homozygous Q84/Q84 mice**

Comparison of phenotypic features in homozygous Q84/Q84 (red triangles, n=11), hemizygous Q84/WT (orange squares, n=8), and wild-type WT/WT (grey circles, n=13) littermates **(A)** Motor impairments in the round balance beam on the first two days of assessments **(B)** Motor impairments in the square balance beam on the first day of assessments **(C)** Minor motor impairments in the accelerating rotarod **(D)** Homozygous mice show a decrease in body weight compared with wild-type and hemizygous littermates. All motor tasks statistics are according to post-hoc Bonferroni corrected estimated marginal means tests of repeated measures ANOVAs. Straight lines: difference between WT/WT and Q84/Q84 mice, capped lines: difference between Q84/WT and Q84/Q84 mice, tick-up lines: difference between WT/WT and Q84/WT mice. Weight statistics according to Bonferroni corrected post-hoc Kruskal Wallis tests. Significance as * $p<0.05$, ** $p<0.01$, *** $p<0.001$.

### Supplementary Figure 1

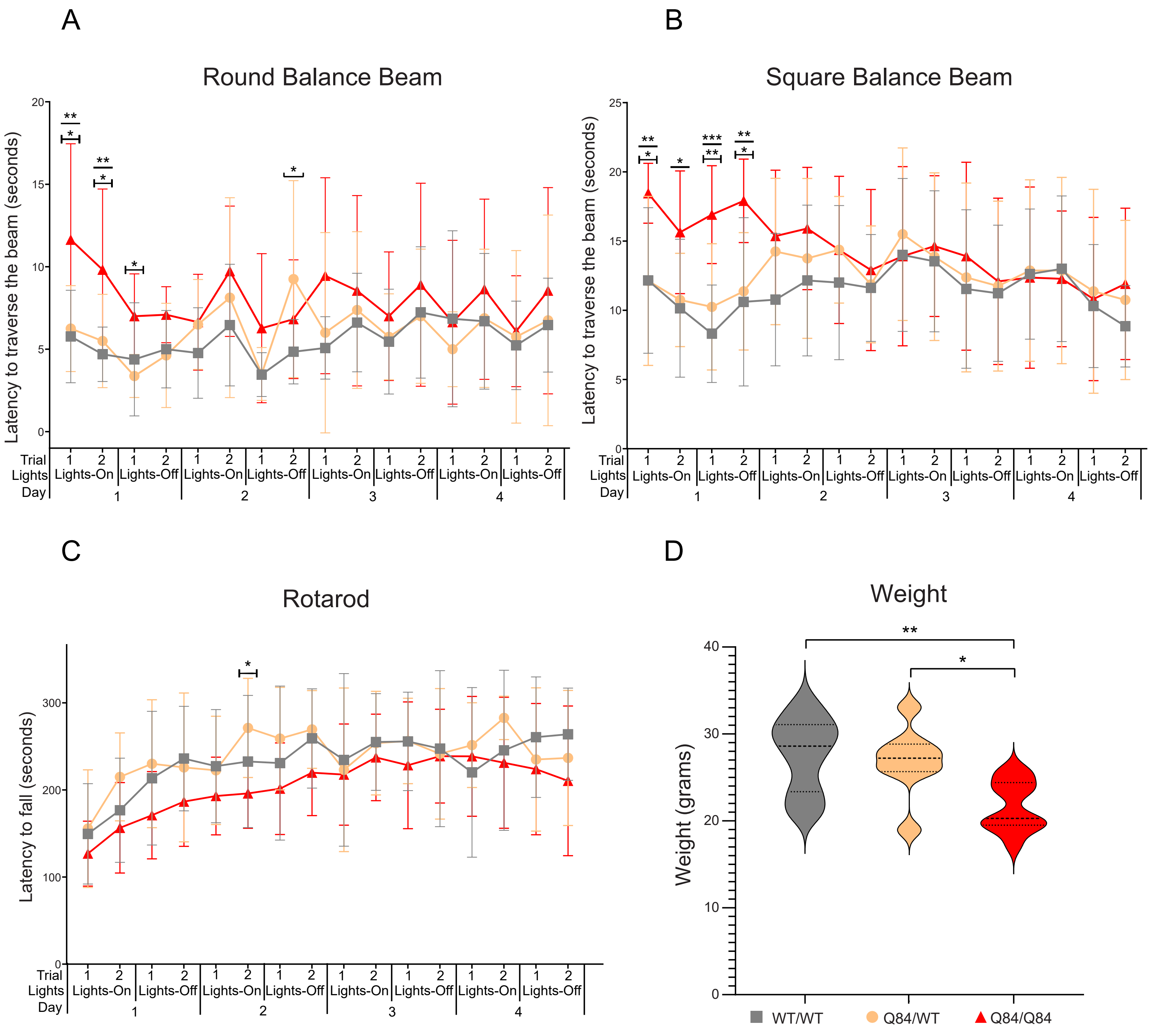
